# An integrated single-cell atlas of the limb skeleton from development through adulthood

**DOI:** 10.1101/2022.03.14.484345

**Authors:** Tim Herpelinck, Liesbeth Ory, Tom Verbraeken, Gabriele Nasello, Mojtaba Barzegari, Johanna Bolander, Frank P. Luyten, Przemko Tylzanowski, Liesbet Geris

## Abstract

The recent growth of single-cell transcriptomics has turned single-cell RNA sequencing (scRNA-seq) into a near-routine experiment. Breakthroughs in improving scalability have led to the creation of organism-wide transcriptomic datasets, aiming to comprehensively profile the cell types and states within an organism throughout its lifecycle. To date, however, the skeleton remains a majorly underrepresented organ system in organism-wide atlases. Considering how the skeleton not only serves as the central framework of the vertebrate body but is also the home of the hematopoietic niche and a central player in major metabolic and homeostatic processes, this presents a major deficit in current reference atlas projects. To address this issue, we integrated ten separate scRNA-seq datasets containing limb skeletal cells and their developmental precursors, generating an atlas of 133 332 cells. This limb skeletal cell atlas describes cells across the mesenchymal lineage from the induction of the limb to the adult bone and encompasses 39 different cell states. Furthermore, expanding the repertoire of available time points and cell types within a single dataset allowed for more complete analyses of cell-cell communication or *in silico* perturbation studies. Taken together, we present a missing piece in the current atlas mapping efforts, which will be of value to researchers in the fields of skeletal biology, hematopoiesis, metabolism and regenerative medicine.

## Introduction

The skeleton is a highly advanced organ with a wide variety of functions, ranging from protection of the internal organs and supporting locomotion to calcium homeostasis, housing the hematopoietic system and serving an endocrine function. In addition, some skeletal tissues possess remarkable regenerative properties, with bone being able to spontaneously regenerate after fracture with minimal scar formation. While complex in its functions, the skeletal system is composed of a surprisingly low number of cell types, suggesting that its functional variety is a trait obtained through extensive intrinsic heterogeneity of cell states allowing advanced regionalized specialization.

Advancement towards ultra-high throughput single-cell RNA sequencing (scRNA-seq) platforms and the concomitant development of computational algorithms required to analyze the data, permit the generation of organism-wide transcriptome maps, resolved in time and space (1,2). If a reference atlas would be available, new datasets could be annotated automatically permitting a fast, data-driven and consistent cell annotation (3–5). Unfortunately, the skeleton is under-represented in most of these atlases, often with insufficiently detailed annotation of the skeletal lineage. Additionally, the skeletal biology research community generated scRNA-seq datasets but focused on very specific cell subpopulations. As a result, we find a significant discrepancy in the transcriptional characterization of the skeletal system. Large atlases contain a wealth of data but are left largely unexplored due to the insufficiently precise annotation while the specialized datasets remain confined to their domain of study. To bridge this gap, we reannotated and integrated specialized skeletal datasets (Supplementary Table 1) into a single framework comprising 133 332 cells (6–15). This reannotation was performed to ensure consistency across all datasets, using the original labels and enhancing them with more detailed annotations through the use of marker genes, resulting in the Limb Skeletal Cell Atlas (LSCA).

Following the generation of the LSCA, we explored its applicability and predictive value. We demonstrated the potential of the LSCA in pseudotemporal trajectory inference and intercellular communication analysis. The predictive capacity of the LSCA was further tested by simulation of Sox9 inactivation and the analysis of its consequences, which were in line with the *in vivo* phenotype. These results support the notion that our LSCA can be used, amongst others, as a reliable reference for the automated annotation of new skeletal scRNA-seq datasets. To facilitate accessibility, all notebooks required to build the atlas are available online.

## Results

### Dataset annotation and integration to produce a limb skeletal cell atlas

To build a comprehensive reference atlas of the limb skeleton, we selected ten publicly available mouse scRNA-seq datasets containing cells from the onset of limb development to mature bone. The individual datasets were manually reannotated based on the original labels and canonical marker gene expression (Supplementary Table 2). A comprehensive analysis was conducted on a total of 133 332 individual cells that met the quality control filters and led to the identification of 39 distinct cell clusters (Fig. 1 a). Twenty-six clusters originated from a mesenchymal lineage and encompassed mesenchyme, chondrocyte, osteoblast/osteocyte or fibroblast cell states. A further four states were associated with the muscle lineage and other clusters included hematopoietic (n=3), endothelial (n=2), epithelial (n=2), ectodermal (n=1) and neuronal (n=1) cell states.

**Figure 1.**
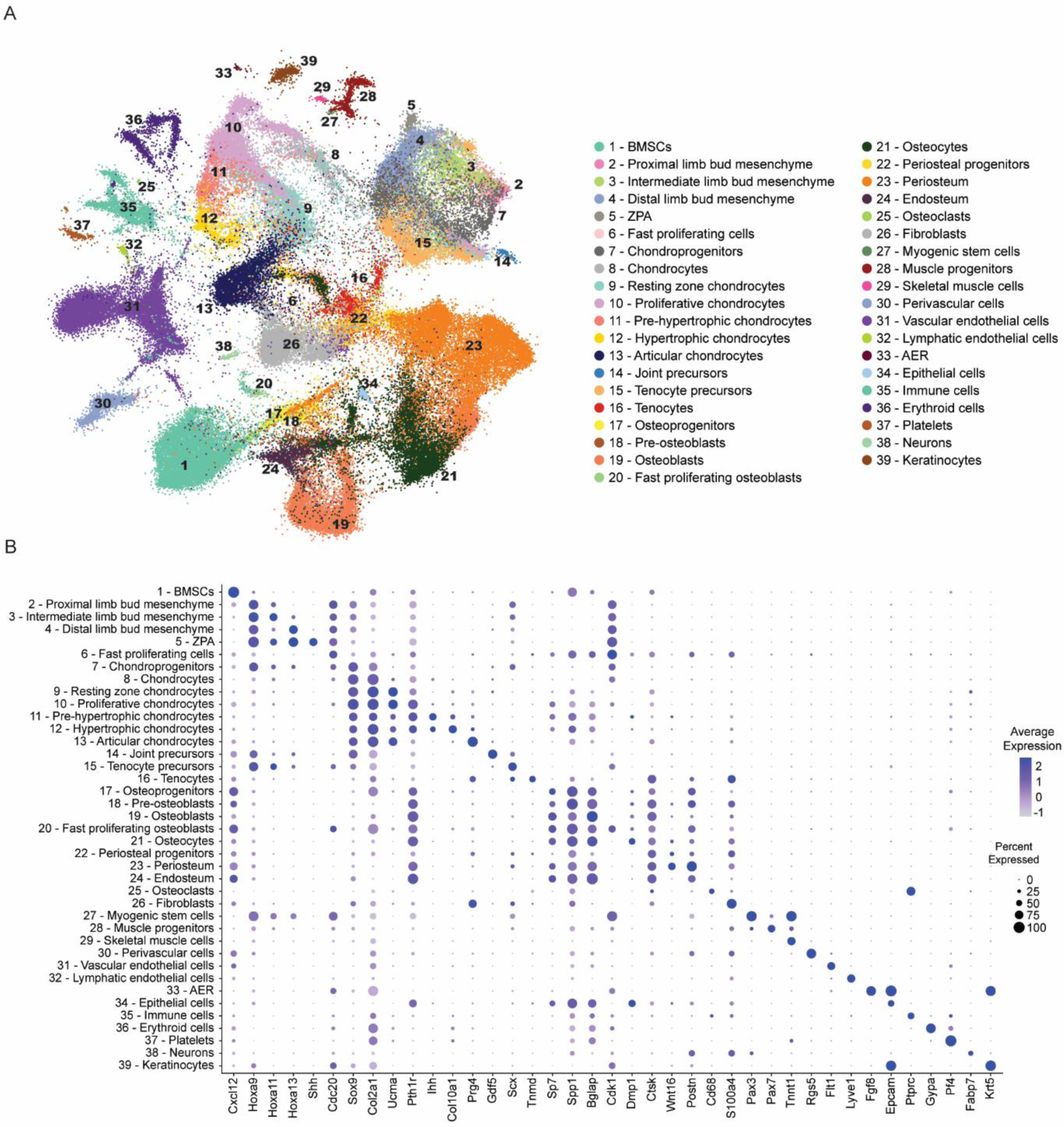
An integrated compendium of skeletal cell types with detailed annotation. a UMAP visualization of the scANVI latent space of 133 332 murine limb mesenchyme- and skeleton-derived cells, colored by annotation. b, Dot plot showing the expression of one selected marker gene per cluster. The color of the dot represents the mean expression level and its size represents the percentage of cells within the cluster in which that gene was detected. BMSCs: Bone Marrow-derived Stem Cells, ZPA: Zone of Polarizing Activity, AER: Apical Ectodermal Ridge.

During single-cell extraction, all spatial data is lost. A pseudospatial context within the cells of the early limb bud was reintroduced utilizing distinct gene expression patterns. Specifically, we used the expression of the Hox gene family to delineate cells along the proximal-distal axis (Hoxa/d9-13) (16). A subset of ectodermal cells situated at the distal tip of the limb bud, called Apical Ectodermal Ridge (AER), is a signaling center associated with the proximal-distal limb growth. It could be identified by the mRNA expression of Fgf8 (17). Cells at the distal-posterior margin of the limb bud exhibited the expression of Shh, designating it as the zone of polarizing activity (ZPA) (18).

Following annotation, we applied both an unsupervised (single-cell Variational Inference; scVI) (19,20) and a semi-supervised (single-cell Annotation using Variational Inference; scANVI) (19,21–24) integration model. The scVI model allowed us to infer the underlying structure of the data without prior labels, while scANVI leveraged both labeled and unlabeled data to improve annotation accuracy. To assess the effectiveness of these integration methods, we employed metrics focused on batch correction and biological conservation (scIB) (19,25) (Supplementary Fig. 2). These metrics helped us evaluate how well the integration methods corrected for batch effects while preserving the biological variability inherent in the data. Notably, scANVI demonstrated the most favorable overall performance, effectively integrating the datasets without introducing cross-laboratory batch effects (Supplementary Fig. 3). This robust performance of scANVI ensured that the integration was cell-based, accurately representing 39 distinct cell states (Fig. 1a-b). This comprehensive integration provides a reliable foundation for downstream analyses and applications.

### Virtual reconstruction of the growth plate

Long bones are formed through the process of endochondral bone formation. Initially, mesenchymal cells condense and then undergo a series of differentiation steps going from resting chondrocytes over proliferating to pre-hypertrophic and hypertrophic chondrocytes. These cells form columnar structures that together make up the growth plate (26). The hypertrophic chondrocytes then either transdifferentiate into osteoblasts or undergo apoptosis and the space is replaced by the osteoblasts producing bone tissue.

As a part of the validation of our atlas, we virtually restored the growth plate. To accomplish that, we used pseudotime analysis to reconstruct the transcriptional trajectories from resting to hypertrophy zone *in silico* (Fig. 2a). Cells were then binned by pseudotime and for each bin, we performed dimensionality reduction by t-distributed stochastic neighborhood embedding (t-SNE). Importantly, we imposed a circle as a boundary condition for each t-SNE. Alignment of these circular projections then recreated the cylindrical shape of the growth plate, while also visualizing transcriptional heterogeneity within the growth plate across bins of pseudotime (Fig. 2b). As such, plotting gene expression along the pseudotime axis combined with expression in pseudospace artificially restores, to a certain extend, the tissue architecture (Fig. 2c).

**Figure 2.**
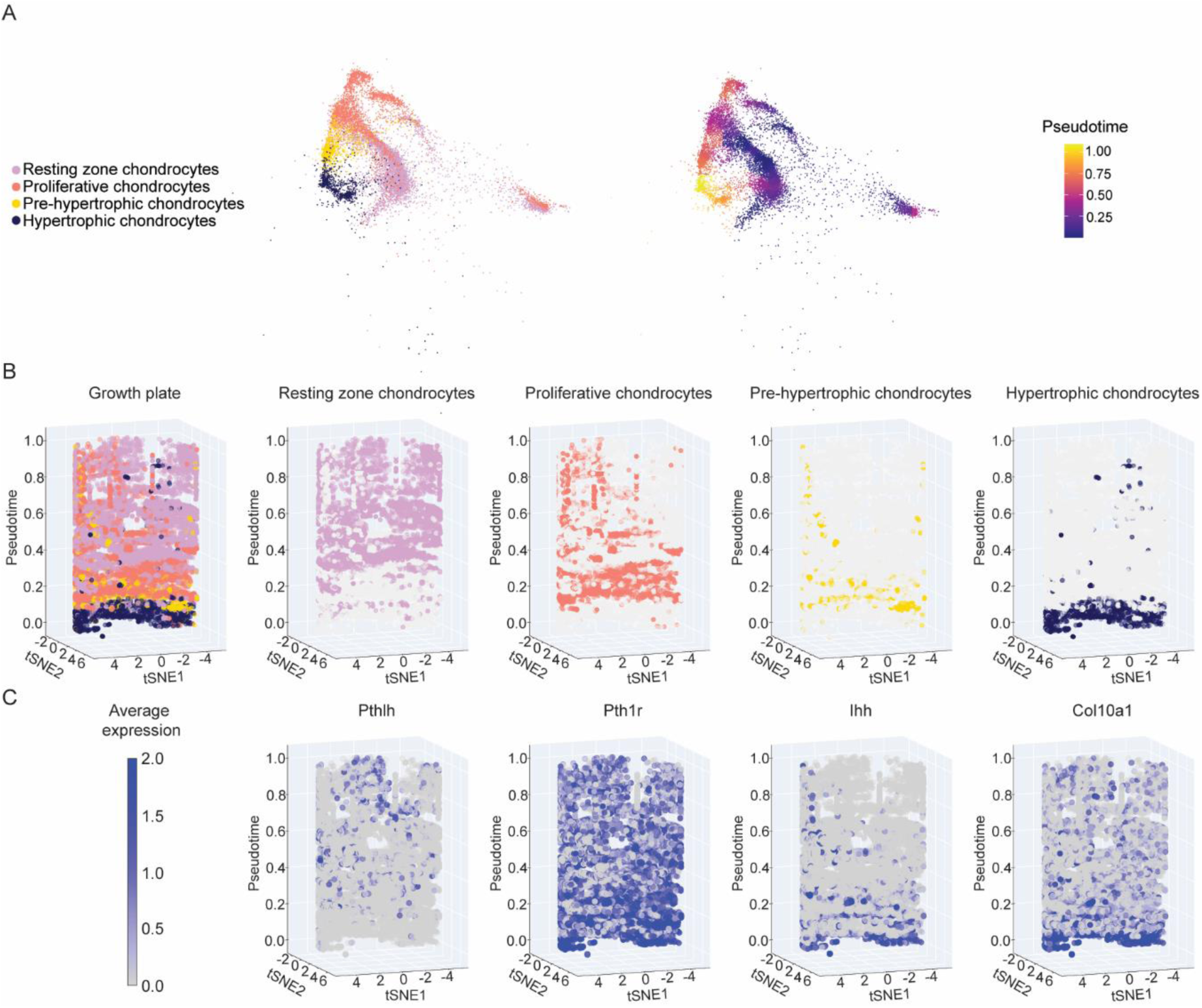
Pseudospatiotemporal reconstruction of the transcriptional dynamics within the growth plate. a, UMAP visualization of the scANVI latent space of the integrated growth plate data subset from the main atlas, colored by annotation (left) and Monocle 3 pseudotime value (right). b, Growth plate chondrocytes were grouped in 50 bins based on similar pseudotime values. t-SNE dimensional reduction was performed on each bin, using a circle with a radius of 20 as the boundary condition for gradient descent, thus recreating the cylindrical shape of the growth plate upon stacking the bins. c, Known marker gene expression for different zones of the growth plate in pseudotime and - space.

### Intercellular communication inference across the skeleton

Intercellular communication driven by ligand-receptor interactions is pivotal for cellular differentiation. Therefore, a wide variety of tools to infer intercellular signaling from scRNA-seq data has been developed (27). Here, we used CellPhoneDB (28) as it considers multimeric receptor complexes when inferring ligand-receptor interactions. We analyzed Bone Morphogenetic Protein (BMP), Growth Differentiation Factor 5 (GDF5) and Sonic Hedgehog (SHH) signaling taking place within the limb bud, the Apical Ectodermal Ridge (AER) and the Zone of Polarizing Activity (ZPA). BMPs are members of the Transforming Growth Factor ligand superfamily and signal through a tetrameric receptor complex. Type I and Type II BMP receptors are expressed across the limb mesenchyme (29,30) and AER (29), which also acts as a source of BMPs (31). However, due to the multimeric character of the receptor complexes resulting in many different combinations, it is challenging to experimentally determine which ligand-receptor pair is predominantly used. Screening with CellPhoneDB narrowed down the number of possible combinations. The analysis revealed that the AER mainly signals to the limb bud through BMP4 and BMP7 (Fig. 3, Supplementary Fig. 4-6) and that they appear to favor binding a receptor complex containing BMR1A. BMP signaling towards the AER from the limb mesenchyme is known to be limited (31), as we confirm in our data. Noteworthy is that the favored ligand-receptor combination is invariable over time. In addition, the AER is predicted to be capable of autocrine signaling, which decreases over time (Fig. 3, Supplementary Fig. 4-6).

**Figure 3.**
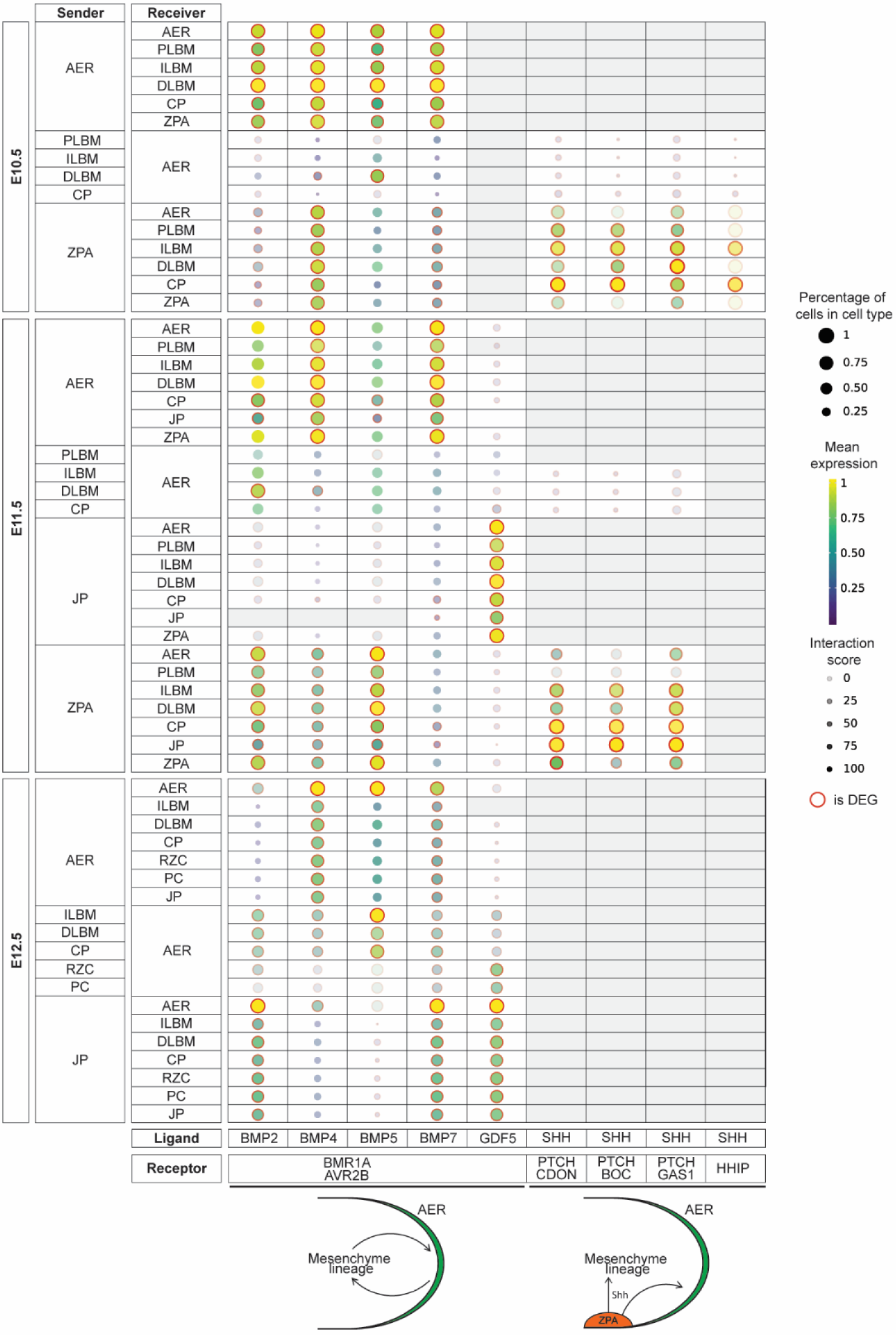
BMP, GDF5 and SHH signaling case study between AER, ZPA and mesenchyme. Dotplot of predicted ligand-receptor complex interactions in developmental timepoints E10.5, E11.5 and E12.5. The sender/receiver cell types are represented on the y-axis, while the x-axis represents the ligand/receptor pairs. The color of the dot represents the mean gene expression, while the size of the dot is proportional to the percentage of cells of each cell type expressing the gene. Translucency represents the interaction specificity and if the gene is differentially expressed an outer red ring is shown. DEG: differentially expressed genes, AER: Apical Ectodermal Ridge, PLBM: Proximal limb bud mesenchyme, ILBM: Intermediate limb bud mesenchyme, DLBM: Distal limb bud mesenchyme, CP: Chondroprogenitors, RZC: Resting zone chondrocytes, PC: Proliferative chondrocytes, PHC: Pre-hypertrophic chondrocytes, JP: Joint precursors, ZPA: Zone of Polarizing Activity.

Previous experiments show that GDF5 binds to different sets of type I receptors. This interaction is essential for the differentiation of digit cartilage and synovial joints (32). Our data shows significant expression from Joint Precursors towards the Mesenchyme and the Chondroprogenitors (Fig. 3).

Antero-posterior outgrowth and proximodistal polarization are interconnected to each other by an SHH epithelial-mesenchymal feedback loop (33). SHH expression decreases as the limb bud grows which explains the decrease in cell-cell communication from E10.5 to E11.5 and no SHH expression in E12.5. SHH signaling is regulated during limb development at many levels and one of them is a negative feedback loop involving binding of SHH to HHIP which acts as a decoy receptor to inhibit HH signaling. As a part of the LSCA validation, we investigated that regulatory loop and showed that this cell-cell communication is mostly limited to intermediate limb bud mesenchyme and chondroprogenitors (Fig. 3).

### Simulation of transcription factor perturbation

Developmental biology was one of the key fields of research for exploring gene regulatory networks (GRNs). Traditionally, this involved a series of experiments where transcription factor activity was altered by gain- or loss-of-function and the resulting *in vivo* effects were analyzed. With the advances in bioinformatics, CellOracle (34) algorithm was designed to study GRN inference from scRNA-seq data. In addition, it allows to explore the *in silico* effects of the perturbation of transcription factor expression on target gene expression. To test if our dataset was amenable to this type of analysis, we decided to compare an *in silico* prediction to published *in vivo* data (35).

Sox9 is a transcription factor essential for maintaining the proliferation of columnar chondrocytes and preventing their premature transdifferentiation into osteoblasts. It also plays a crucial role in inducing chondrocyte hypertrophy, making it indispensable for the proper formation and maintenance of functional growth plates. Inactivation of Sox9 leads to a shortening of the columnar and hypertrophic zones in the growth plate and accelerates ossification due to premature pre-hypertrophy and matrix mineralization (11,36). To assess the usefulness of our atlas for functional studies, we applied the CellOracle algorithm to create a virtual Sox9 knockout and compared our *in silico* predictions to data from an *in vivo* knockout study (Fig. 4a,b). We identified regions within the dataset that are developmentally accessible or inaccessible depending on Sox9’s regulatory influence. Accessible cell states are those that cells can develop into under the current regulatory conditions, while inaccessible states cannot be reached due to the absence or inactivation of essential transcription factors.

**Figure 4.**
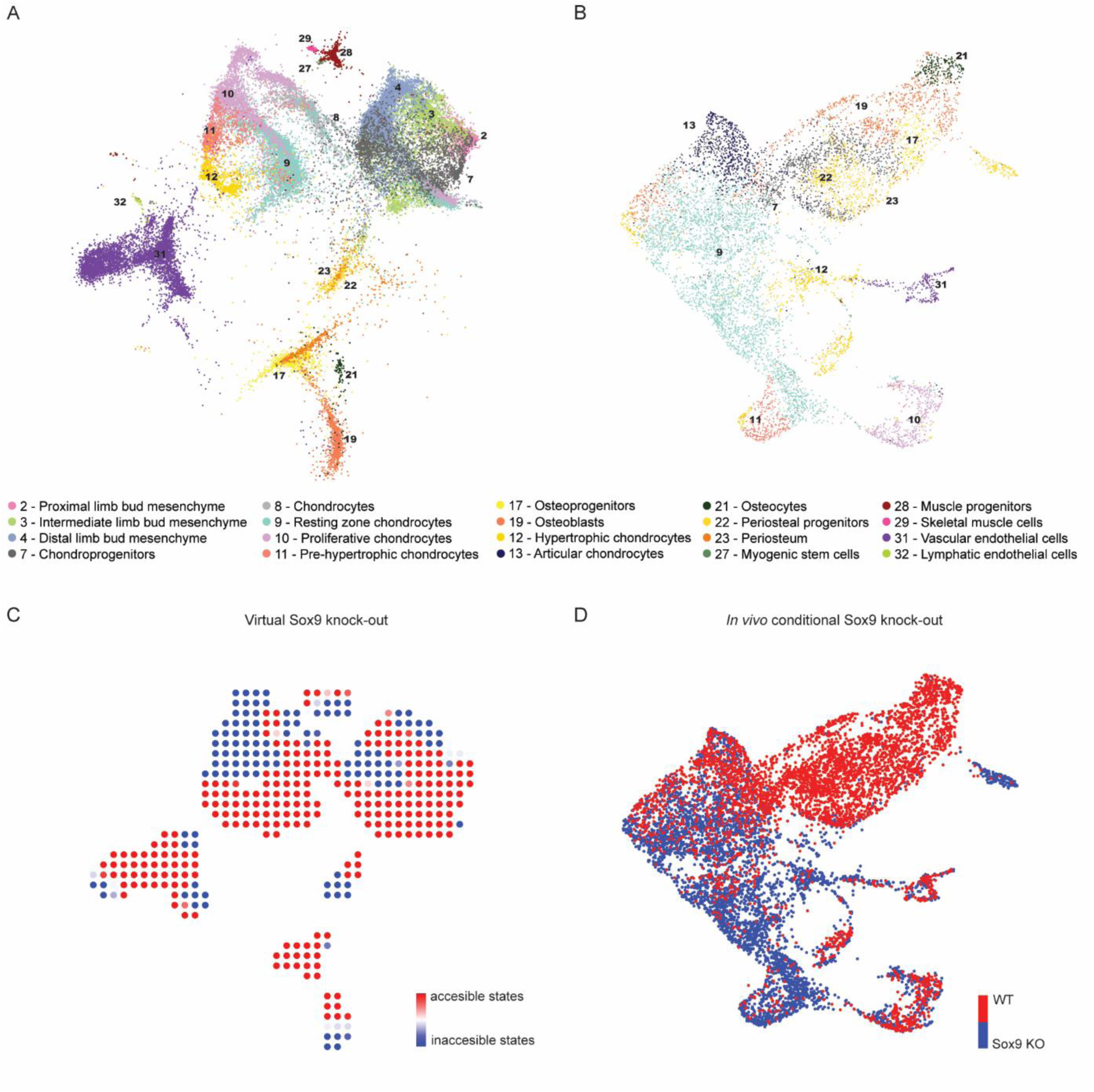
Sox9 knockout (KO) *in silico* predictions compared to *in vivo* data. a, Reference annotation of the subsetted atlas used for knockout analysis. b, a UMAP plot integrating WT and Sox9 KO *in vivo* data. c, Inner vector product of cell trajectories following *in silico* Sox9 perturbation and the pseudotime gradient from unperturbed conditions. Red regions indicate accessible cell states, while blue regions represent inaccessible cell states. d, UMAP of integrated Sox9 KO and WT *in vivo* data, colored by condition. Red corresponds to Sox9 KO data, while blue represents WT data. WT: Wild-type, KO: knockout.

The simulations show that upon removal of Sox9, most mesoderm-derived cells fail to contribute to cartilage formation, while myogenesis, osteogenesis, and endothelial differentiation remain largely unaffected (Fig. 4c, Supplementary Fig. 7). These results align with *in vivo* findings, where Sox9 inactivation prevents mesodermal cells from differentiating into chondrocytes, leading to an accumulation of cartilage precursors and osteogenic cells (Fig. 4d). For example, in the virtual knockout model, cell type 11—pre-hypertrophic chondrocytes—is represented as a blue region, indicating that mesenchymal cells cannot access this cell state. This observation is consistent with *in vivo* data, where this cell type is exclusively present in the wild-type (blue) and absent in the knockout.

A limitation of the computational approach to create a virtual knockout is that it accounts only for developmental flow and not lineage dependency. As a result, it will mark a region as an inaccessible cell state if its development depends on Sox9, however, it will mark regions relying on this (inaccessible) cell type as accessible states. For instance, cell type 12—hypertrophic chondrocytes—is shown as a red region in the virtual knockout, indicating that this cell state is predicted to be present in Sox9-perturbed situations. However, *in vivo* data show that this cell type is absent in the knockout (blue) because it originates from pre-hypertrophic cells.

## Discussion

We have generated a Limb Skeletal Cell Atlas representing a manually curated compendium of 133 332 cells across ten datasets and explored its applicability in both data- and hypothesis-driven analyses. First, we presented the unsupervised construction of a spatiotemporal map of the growth plate, which correctly recapitulates the molecular cascades of growth plate development, by imposing boundary conditions on the gradient descent of t-SNE dimensionality reduction. Similar results have been obtained previously by warping the principal component space (37).

Next, analysis of intercellular communication by BMP signaling was able to shed light on a longstanding question in the field of limb development. It is known that BMP signaling is required for AER regression, but not whether the AER, the mesenchyme or both act as the source of those BMPs (31). Based on CellPhoneDB’s cell-cell communication inference, considering the subunit architecture of both ligands and heteromeric receptors (27,28), our case study suggests the AER to be the dominant source. This finding does, however, require further *in vivo* validation.

Finally, we demonstrated that the *in silico* inactivation of Sox9 corroborates *in vivo* results confirming the inability of Sox9-negative cells to differentiate into chondrocytes (11,38). Given that we can assess the developmental outcome of computationally simulated transcription factor knockouts, the LSCA represents a valuable resource for both limb developmental biologists and skeletal biology researchers.

The current release is restricted to the healthy murine skeleton. While our study provides valuable insights, it is important to acknowledge certain limitations. Specifically, there are gaps in the limb sites and stages that were sampled, as well as differences in genetic background, sex, and the enrichment of sub-populations. These factors may influence the census and its potential downstream applications. However, these limitations also highlight opportunities for future research to build upon our findings, addressing these gaps and further enhancing our understanding of the subject. We will continuously update and extend the LSCA, as data availability permits, toward the entire limb skeleton in health and disease. The goal of this study was to provide an initial reference of the musculoskeletal system of the limb. In future work, this first version will serve as the basis of a collaborative effort to complete the atlas. All data, analyses and code to replicate the figures are freely available. Our web portal is designed to be intuitive and to allow browsing of the data at a glance. In short, the Limb Skeletal Cell Atlas provides a framework with characterization of most known cell populations in the limb skeleton and represents a foundation for future studies in a wide variety of disciplines.

## Materials and Methods

### Data preprocessing

Publicly available datasets that were not aligned, were first mapped with Cell Ranger (39). Initial Quality control of the raw count matrices was then performed for each dataset using Scater (40). Filtering was performed based on the library size and number of expressed genes per cell. Cells more than three standard deviations away from the median of either metric were filtered out. Subsequently, cells where the mitochondrial fraction of reads exceeded a proportion higher than three standard deviations from the median mitochondrial fraction were removed. Finally, low-abundance genes (expression lower than 1E-3) were filtered out and duplicate rows were removed if present.

### Clustering and dimensionality reduction

All datasets were analyzed individually prior to integration. The filtered count matrices were imported into Seurat v4 (4). Normalization was performed using the LogNormalize parameter and a scale factor of 1E4. Subsequently, the data was centered and scaled using all genes followed by a calculation of principal components (PCs) using the top 2000 highly variable genes selected by the “vst” method. The optimal number of PCs to construct the Shared Nearest Neighbor (SNN) graph was visually determined based on the elbow plot and varied for each dataset. Clustering was performed using the Louvain algorithm. The resolution was adapted to the individual dataset, where we defined the optimal number of clusters as the maximum number of cell states that could confidently be labeled based on marker gene expression. These marker genes were obtained from the FindMarkers function using the default settings.

### Integration with scvi-tools

The integrated atlas was constructed using scVI (20) and scANVI (21–24). For preprocessing the datasets, we filtered for the 5000 most variable genes. First, we used scVI as an unsupervised tool to find common axes of variation between the datasets. The parameters were used as described in the scvi-tools (19) tutorial of ‘*Atlas-level integration of lung data*’. Next, to compute a more accurate integration result, we used scANVI as a semi-supervised tool to integrate the data. scANVI used the trained scVI model to initialize the integration.

### Integration metrics

To calculate integration metrics, the scIB-metrics package was used (25), which contains scalable implementations of the metrics used in the scIB benchmarking suite (41–44).

### Virtual growth plate reconstruction

We first subsetted the atlas for growth plate chondrocytes: Resting zone chondrocytes, Proliferative chondrocytes, Pre-hypertrophic chondrocytes and Hypertrophic chondrocytes. This subset within the latent space of the scANVI integration was then passed to Monocle 3 (45). Clustering was performed on the resulting CellDataSet (CDS) object using the default parameters. The trajectory graph was learned on the monocle-derived clusters by calling learn_graph. Roots were manually chosen with order_cells. The resulting pseudotime was added to the metadata of the growth plate subset within the latent space of scANVI and used as input for the growth plate reconstruction. Cells were binned by pseudotime using the cut function from-pandas (46,47). Upon each bin, we performed dimensionality reduction by t-distributed stochastic neighborhood embedding (t-SNE). We imposed a circle as a boundary condition on the gradient descent function of each t-SNE. With the concatenation function, we were able to stack the circular projections in an ordered way to recreate the cylindrical shape of the growth plate.

### Cell-cell interaction prediction

Prediction of cell-cell communication by ligand-receptor interactions between cell types was performed using CellPhoneDB v5 (28). The atlas was subsetted for each developmental time point before downstream cell-cell communication analysis. For each subset, the translate function was used to humanize the data and normalized. Differentially expressed genes were computed for each cell type and used as input for ligand-receptor pair calculations.

### Knockout simulation

For *in silico* knockout experiments in CellOracle (34) the atlas was subsetted to only include time points E10.5-P21 and relevant cell types (Periosteal progenitors, Periosteum, Endosteum, Resting zone chondrocytes, Proliferative chondrocytes, Pre-hypertrophic chondrocytes, Hypertrophic chondrocytes, Chondrocytes, Chondroprogenitors, Osteoprogenitors, Pre-osteoblasts, Osteoblasts, Osteocytes, Vascular endothelial cells, Lymphatic endothelial cells, Proximal limb bud mesenchyme, Intermediate limb bud mesenchyme, Distal limb bud mesenchyme, Myogenic stem cells, Muscle progenitors and Skeletal muscle cell). This subset within the latent space of the scANVI integration of the atlas was then passed to Monocle 3 (45). Clustering was performed on the resulting CellDataSet (CDS) object using the default parameters. The trajectory graph was learned on the monocle-derived clusters by calling learn graph. Roots were manually chosen with order cells. The resulting pseudotime was added to the metadata of the knockout subset within the latent space of scANVI and used as input for the virtual knockout. To reduce the amount of computational time and resources required by a large dataset, 20,000 cells were randomly selected and only highly variable genes (n=5000) were included. For gene regulatory network (GRN) inference, we used the built-in base GRN made from the mouse sci-ATAC-seq atlas (48) Following k nearest neighbors (KNN) imputation based on the first 27 PCs, GRNs were imputed for each cluster. To simulate the knockout of a transcription factor, its expression was set to 0. After this knockout, GRN inference was performed again. Signal perturbation propagation and transition probabilities were calculated using the standard settings. Visualization of the pseudotime gradient, simulation vector field and their inner product was performed as described in the CellOracle online documentation.

### *In vivo* Sox9 knockout and WT data analysis and integration

Both wild type (WT) and Sox9 knockout (KO) datasets were preprocessed separately to ensure high-quality data for subsequent analyses. The preprocessing steps included quality control measures to remove low-quality cells, as described in section 4.1. Following preprocessing, cells from both WT and Sox9 KO datasets were clustered and labeled independently. Clustering was conducted to identify distinct cell populations within each condition, and labels were assigned based on known cell markers, as described in section 4.2.

To integrate the WT and Sox9 KO datasets, the Harmony package (41) was employed. Harmony facilitates the integration of multiple datasets by correcting batch effects and aligning data across different conditions. This integration enabled a comprehensive comparison of cell populations and gene expression profiles between WT and Sox9 KO conditions.

## Data and Code Availability Statement

The datasets analyzed for this study can be found in Supplementary Table 1. The Skeletal Cell Atlas can be downloaded or interactively explored at www.skeletalcellatlas.org. All code used to perform the analyses and notebooks to generate the figures is available at https://github.com/TElabSBE/LimbSkeletalCellAtlas.

## Supplementary Data Figure Legends

**Supplementary Figure 1.**
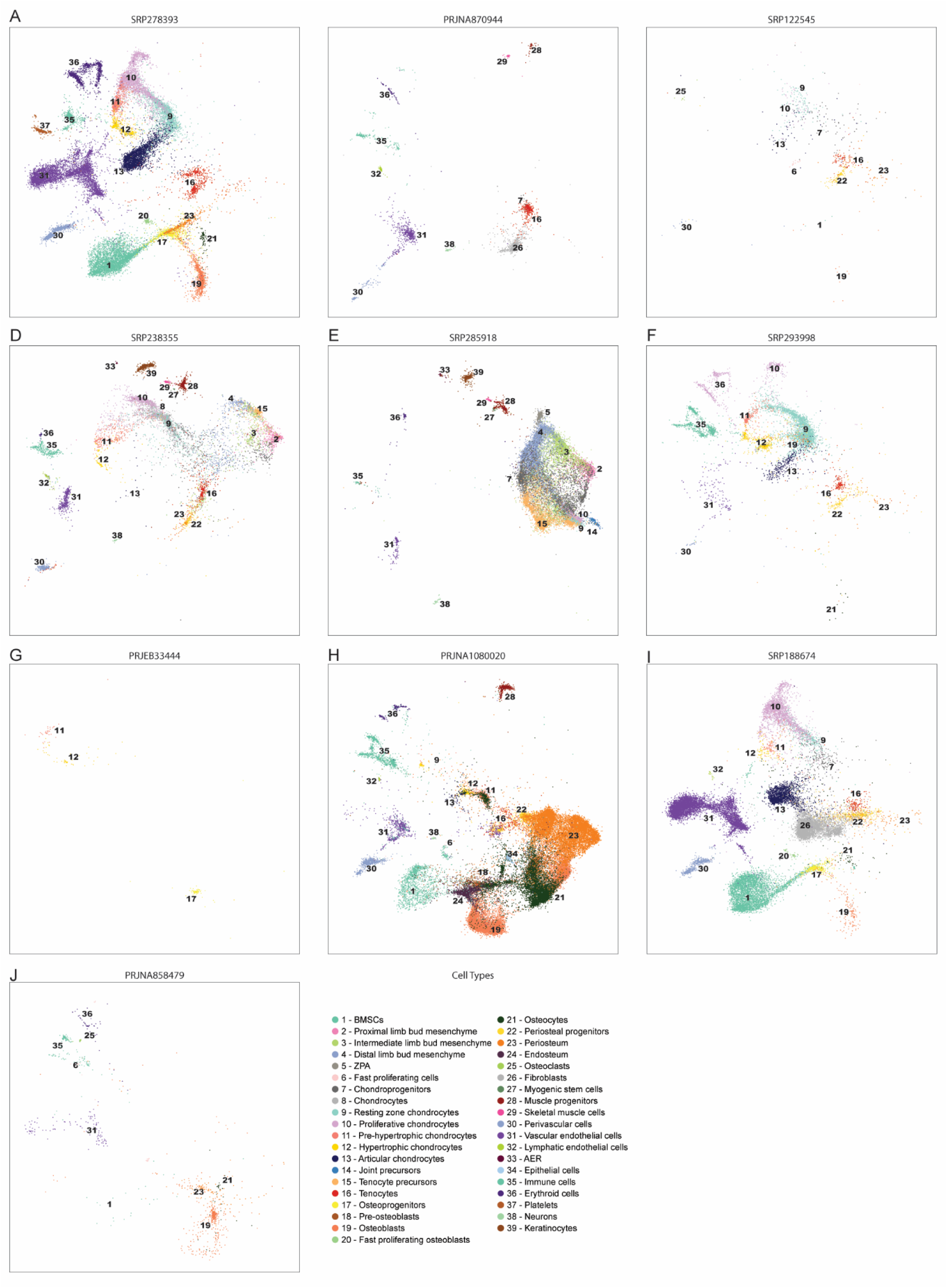
Individual datasets after reannotation. a-j, UMAP visualization of the scANVI latent space colored by cell type for the separate datasets with database ID: a, 34260921 b, 36175067, 37873464 c, 30250253 d, 31874220 e, 33297480 f, 33597301 g, 31543445 h, 38479598 i, 31130381 j, 36777346

**Supplementary Figure 2.**
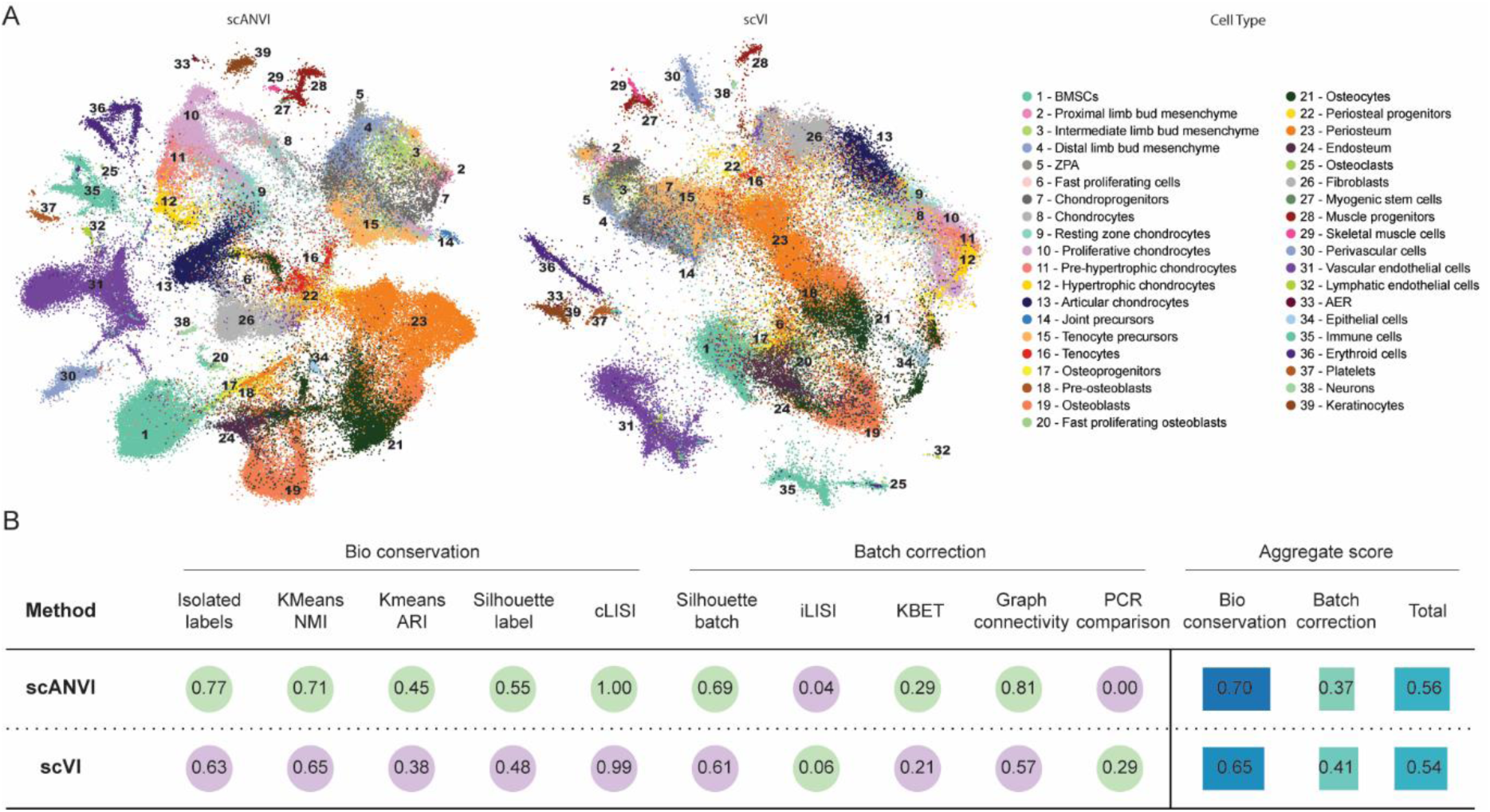
Assessment of integration method. a, UMAP visualization of the scANVI latent space (left) and the scVI latent space (right). b, Results of the metrics calculated with scIB. Bio conservation was captured with the use of classical label conservation metrics, which assess local neighborhoods (graph cLISI, extended from cLISI), global cluster matching (Adjusted Rand Index: ARI, normalized mutual information: NMI and a metric evaluating rare cell identity annotations (isolated label scores). Batch correction scores were measured via the k-nearest-neighbor batch effect test (kBET), k-nearest-neighbor (kNN) graph connectivity and the average silhouette width (ASW) across batches. Independently of cell identity labels, batch removal is measured using the graph integration local inverse Simpson’s Index (graph iLISI, extended from iLISI) and PCA regression. Aggregate scores represent the mean score of the Bio conservation and Batch correction metrics.

**Supplementary Figure 3.**
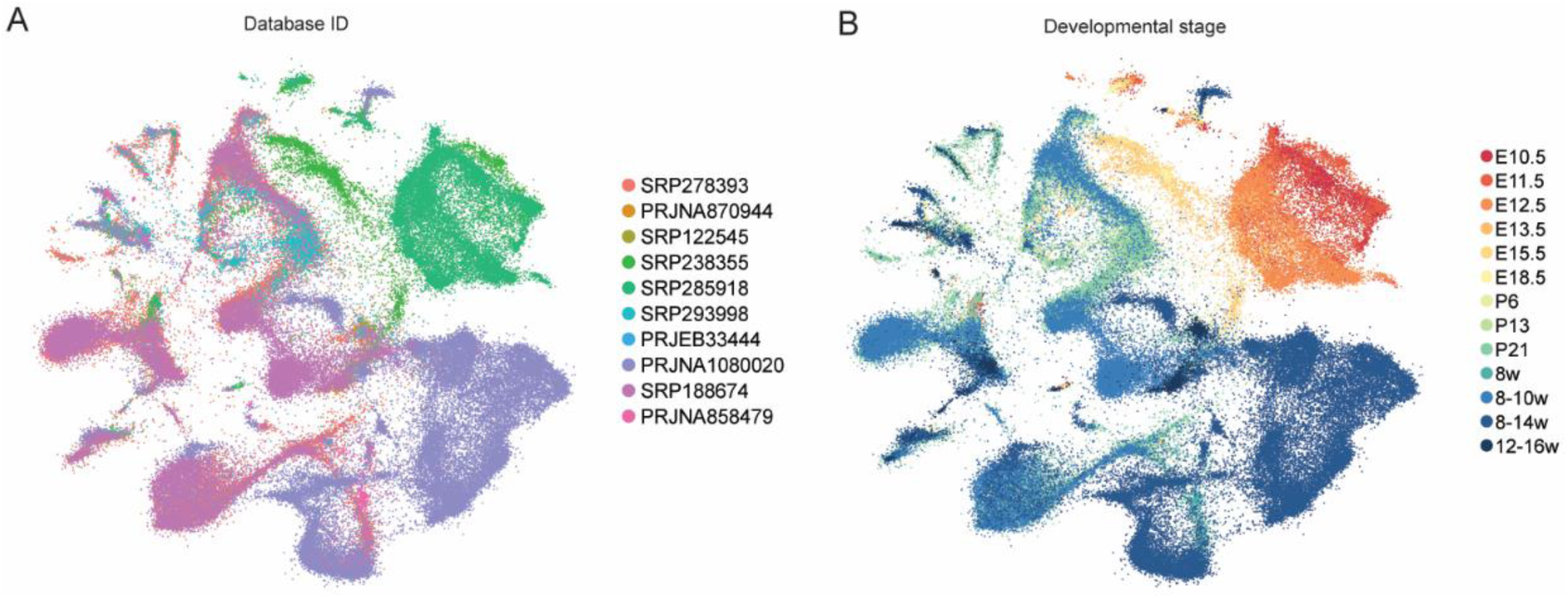
Assessment of batch effects influencing integration. a-b, UMAP visualization of the scANVI latent space colored by study (a) and developmental stage or age (b).

**Supplementary Figure 4.**
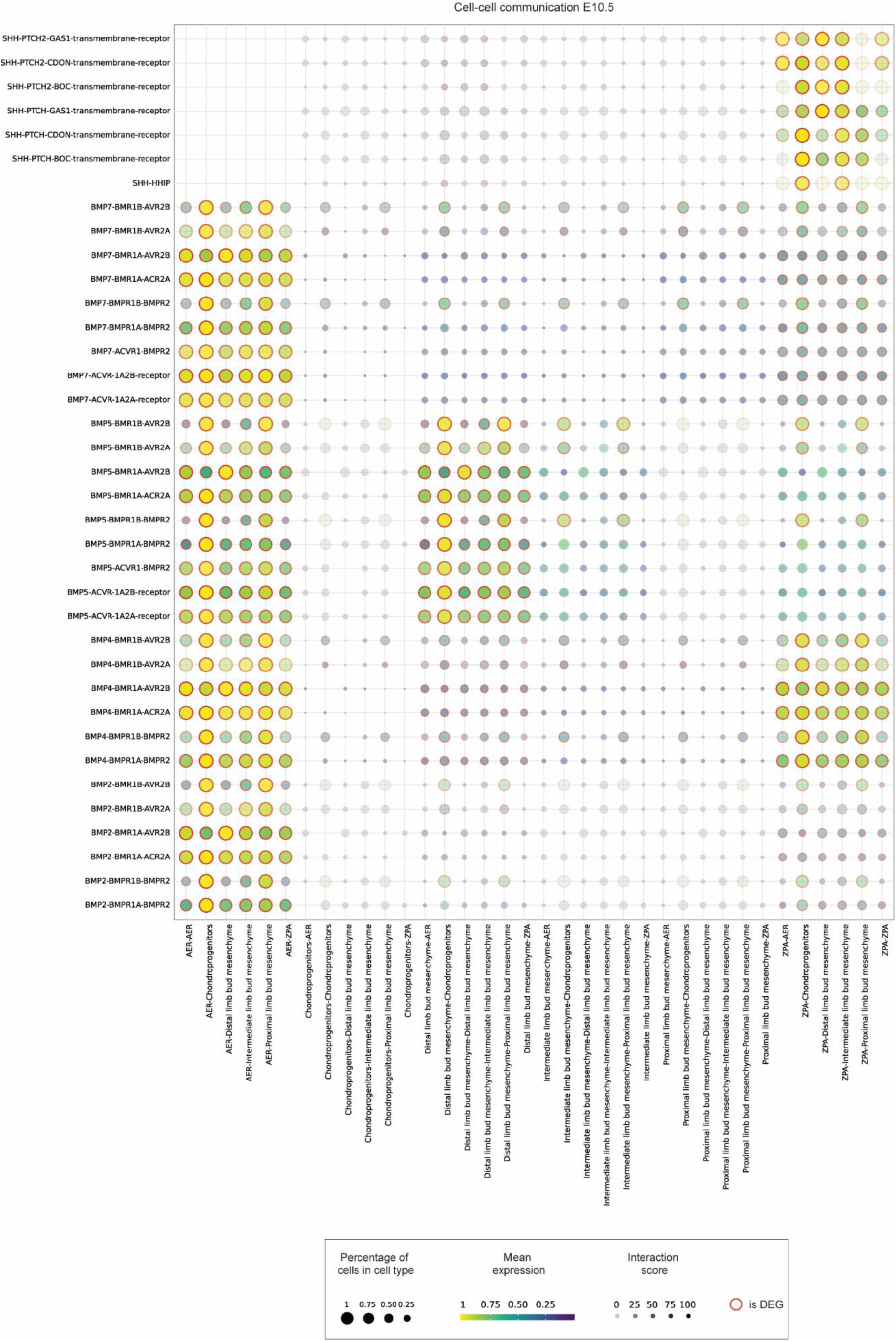
BMP, GDF5 and SHH signaling case study between AER, ZPA and mesenchyme for developmental timepoint E10.5. Dotplot of predicted ligand-receptor complex interactions in developmental timepoints E10.5. The sender/receiver cell types are represented on the y-axis, while the x-axis represents the ligand/receptor pairs. The color of the dot represents the mean gene expression, while the size of the dot is proportional to the percentage of cells of each cell type expressing the gene. Translucency represents the interaction specificity and if the gene is differentially expressed an outer red ring is shown. DEG: differentially expressed genes, AER: Apical Ectodermal Ridge, PLBM: Proximal limb bud mesenchyme, ILBM: Intermediate limb bud mesenchyme, DLBM: Distal limb bud mesenchyme, CP: Chondroprogenitors, RZC: Resting zone chondrocytes, PC: Proliferative chondrocytes, PHC: Pre-hypertrophic chondrocytes, ZPA: Zone of Polarizing Activity.

**Supplementary Figure 5.**
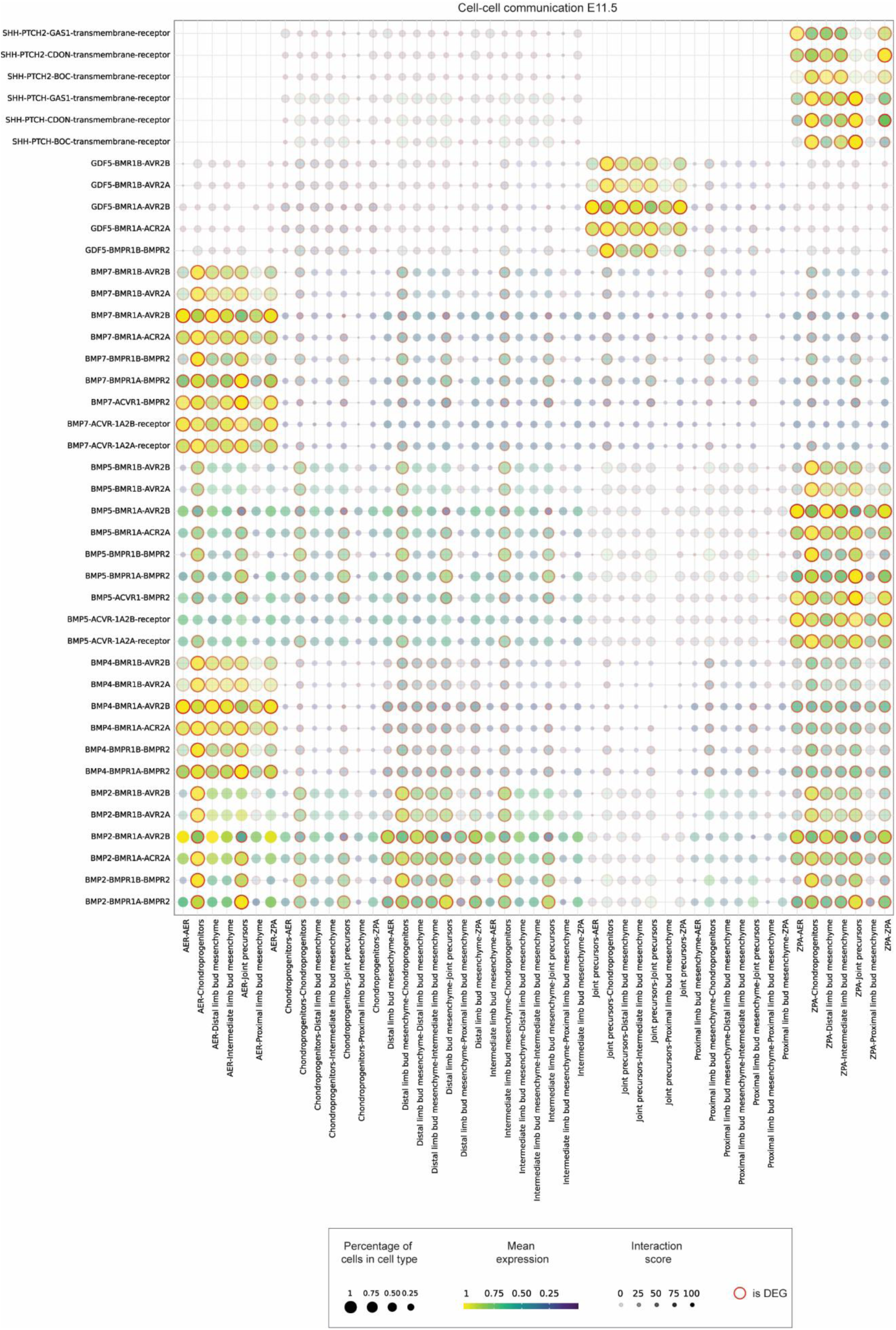
BMP, GDF5 and SHH signaling case study between AER, ZPA and mesenchyme for developmental timepoint E11.5. Dotplot of predicted ligand-receptor complex interactions in developmental timepoints E11.5. The sender/receiver cell types are represented on the y-axis, while the x-axis represents the ligand/receptor pairs. The color of the dot represents the mean gene expression, while the size of the dot is proportional to the percentage of cells of each cell type expressing the gene. Translucency represents the interaction specificity and if the gene is differentially expressed an outer red ring is shown. DEG: differentially expressed genes, AER: Apical Ectodermal Ridge, PLBM: Proximal limb bud mesenchyme, ILBM: Intermediate limb bud mesenchyme, DLBM: Distal limb bud mesenchyme, CP: Chondroprogenitors, RZC: Resting zone chondrocytes, PC: Proliferative chondrocytes, PHC: Pre-hypertrophic chondrocytes, JP: Joint precursors.

**Supplementary Figure 6.**
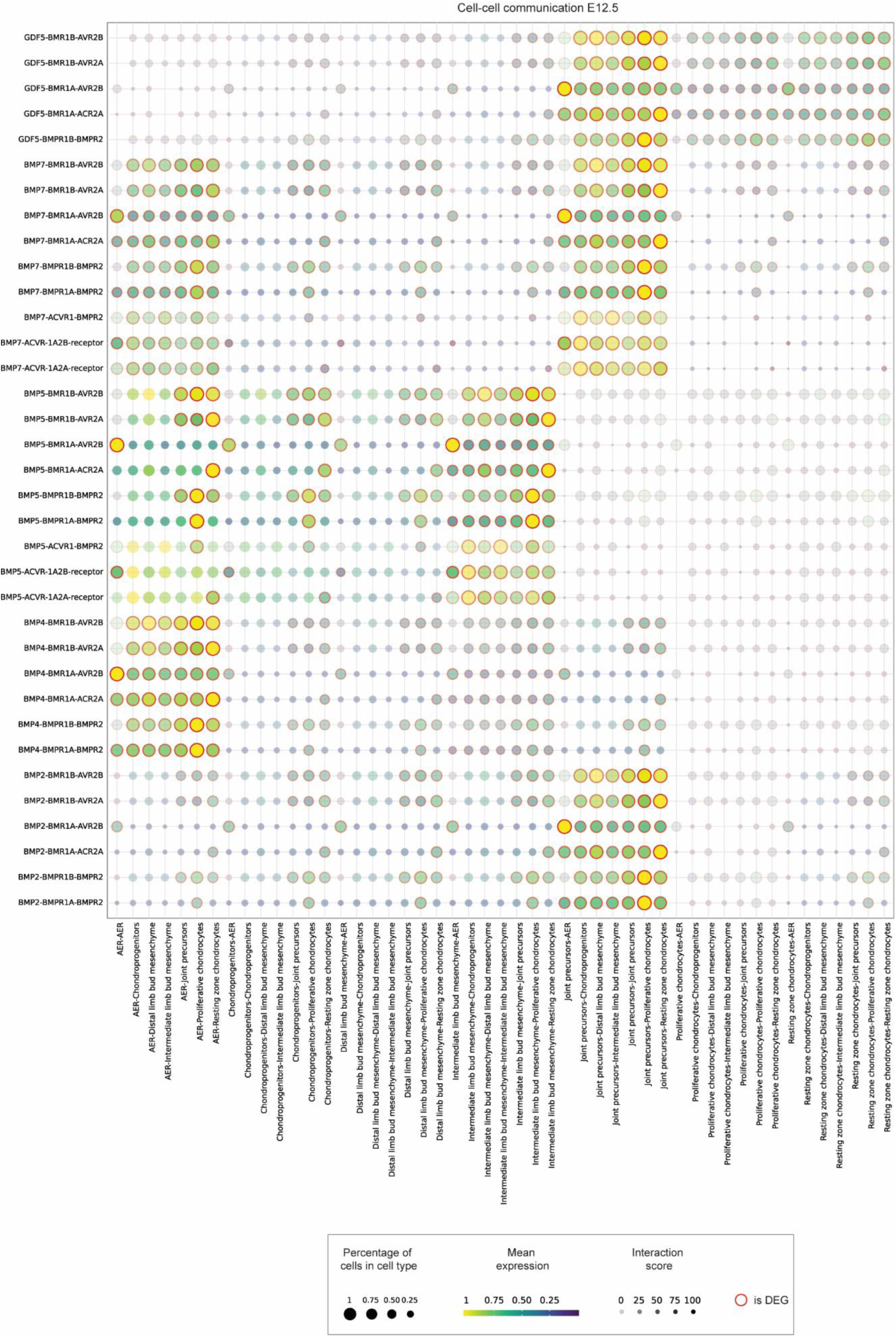
BMP, GDF5 and SHH signaling between AER and mesenchyme for developmental timepoint E12.5. Dotplot of predicted ligand-receptor complex interactions in developmental timepoints E12.5. The sender/receiver cell types are represented on the y-axis, while the x-axis represents the ligand/receptor pairs. The color of the dot represents the mean gene expression, while the size of the dot is proportional to the percentage of cells of each cell type expressing the gene. Translucency represents the interaction specificity and if the gene is differentially expressed an outer red ring is shown. DEG: differentially expressed genes, AER: Apical Ectodermal Ridge, PLBM: Proximal limb bud mesenchyme, ILBM: Intermediate limb bud mesenchyme, DLBM: Distal limb bud mesenchyme, CP: Chondroprogenitors, RZC: Resting zone chondrocytes, PC: Proliferative chondrocytes, PHC: Pre-hypertrophic chondrocytes, JP: Joint precursors, ZPA: Zone of Polarizing Activity.

**Supplementary Figure 7.**
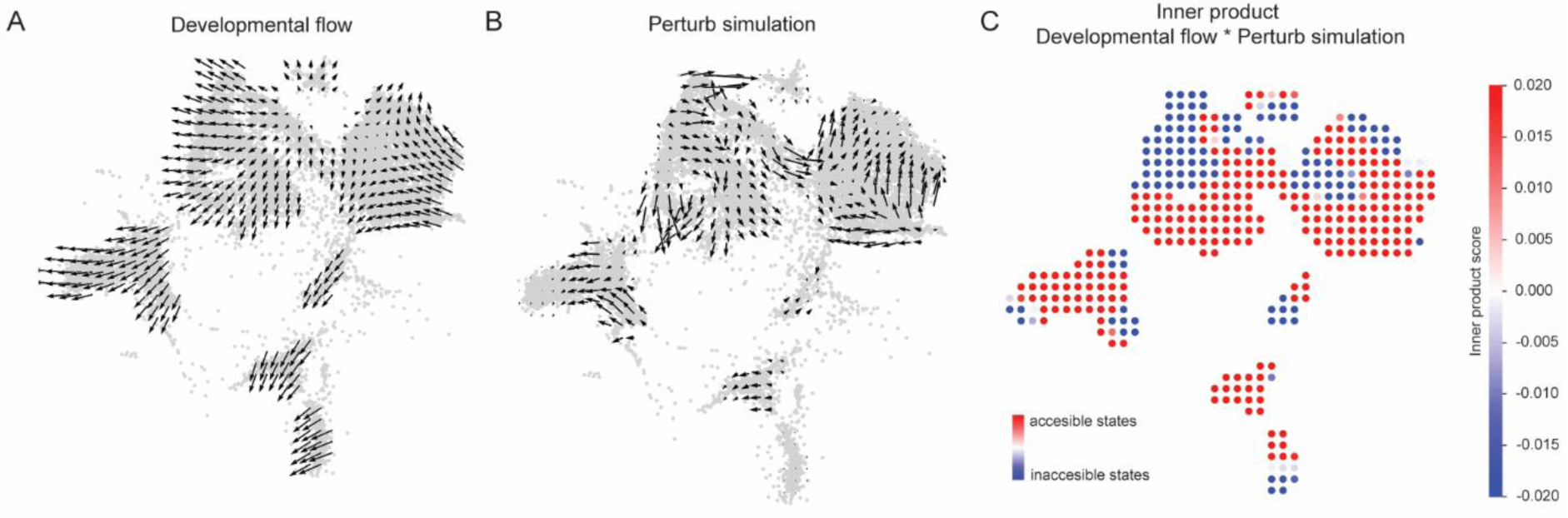
Sox9 knockout (KO*) in silico* predictions. a, Vector field graph indicating cell trajectories after Sox9 *in silico* knockout. b, Gradient of pseudotime in the absence of perturbation. c, Inner vector product of cell trajectories following *in silico* Sox9 perturbation and the pseudotime gradient from an unperturbed condition.

## Supplementary Table Legends

**Supplementary Table 1:** scRNA-seq datasets used to construct the Limb Skeletal Cell Atlas.

| Study (Database ID) | Study (PMID) | Study (First, Last Author) | Technology | Developmental stage |
| --- | --- | --- | --- | --- |
| SRP278393 | 34260921 | Sivaraj, Adams | 10x | P21 |
| PRJNA870944 | 36175067,<br>37873464 | Maerz, Farrell | 10x | 12-16w |
| SRP122545 | 30250253 | Debnath, Landau & Greenblatt | CEL-seq2 | P7 |
| SRP238355 | 31874220 | Kelly, Guilak | 10x | E11.5, E13.5, E15.5, E18.5 |
| SRP285918 | 33297480 | Desanlis, Kmita | 10x | E10.5, E11.5, E12.5 |
| SRP293998 | 33597301 | Lefebvre | 10x | P13 |
| PRJEB33444 | 31543445 | Bohm, Maes | Smart-seq2 | E15.5 |
| PRJNA1080020 | 38479598 | OBrien | 10x | 8-14w |
| SRP188674 | 31130381 | Baryawno, Regev & Scadden | 10x | 8-10w |
| PRJNA858479 | 36777346 | White | 10x | 8w |

**Supplementary Table 2:**
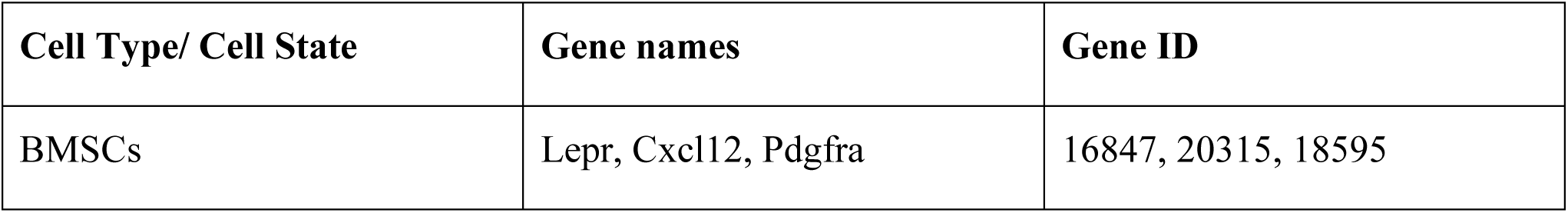
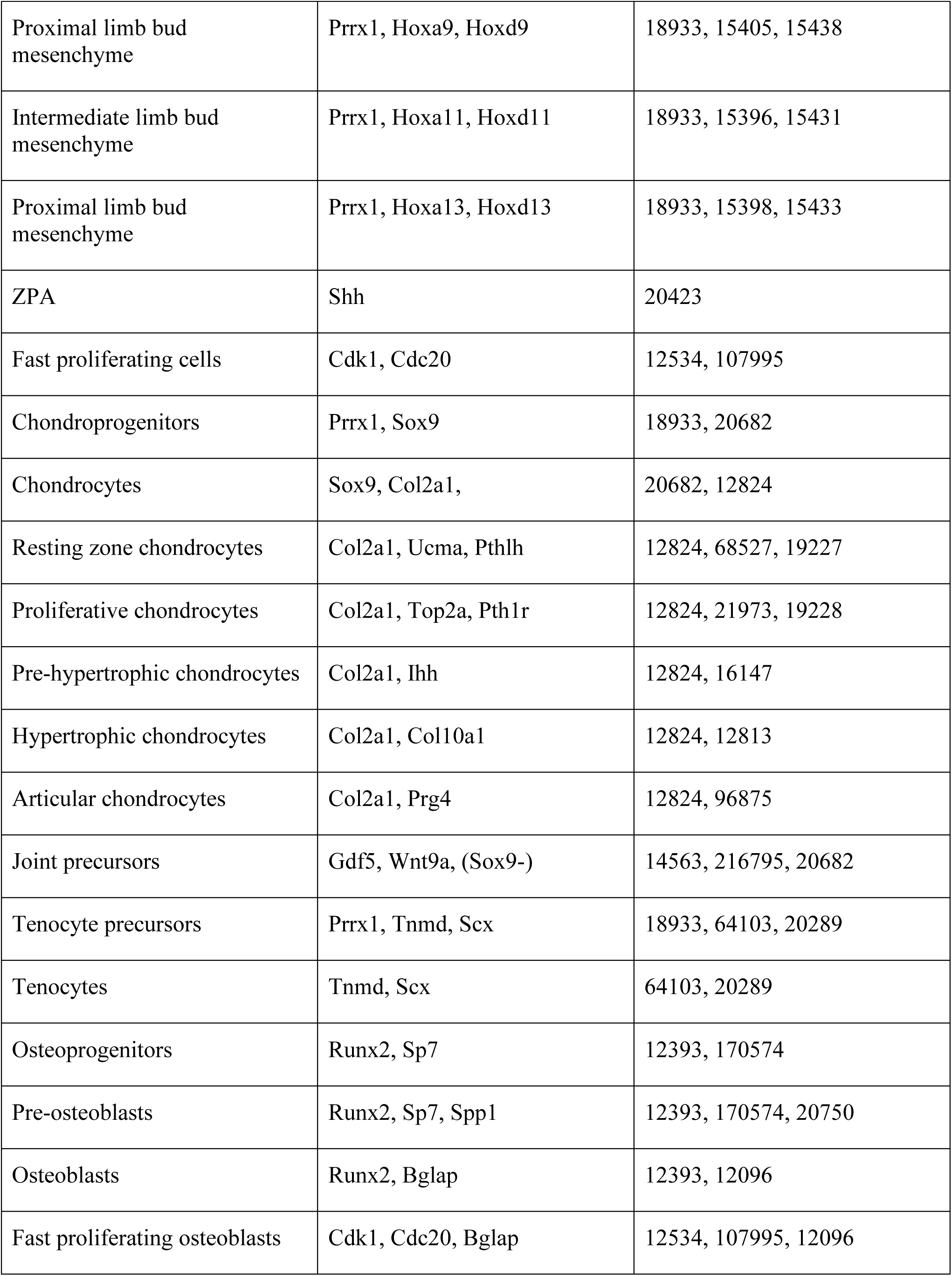
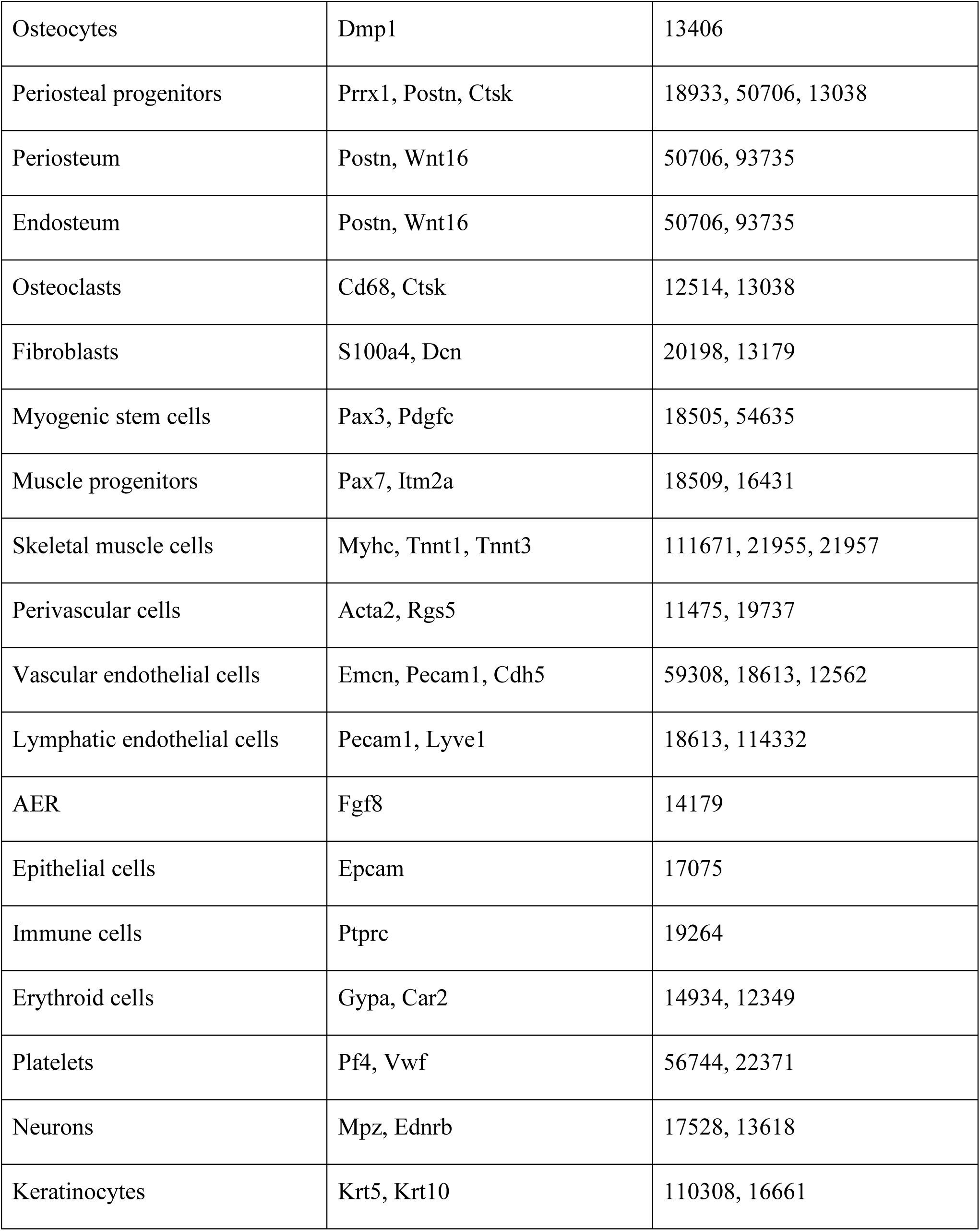
List of genes used to annotate clusters in the Limb Skeletal Cell Atlas.

## Author Contributions

TH, JB, FL, PT and LG conceptualized the study; TH, PT and LG designed the study; TH, LO and TV provided adapted code for the analysis of the datasets and downstream analysis of the LSCA; TH, LO and TV contributed to the analysis of the datasets and the LSCA; GN provided a docker work environment; MB adapted and optimized the code to create an interactive web application, TH and LO wrote the first draft of the manuscript and all the other authors contributed to the article. All authors approved the submitted version.

## Acknowledgments

We would like to thank Inge Van Hoven for assistance and Ronny Moreas for technical support in the deployment of the web app. Research was funded by the Research Foundation Flanders (FWO Vlaanderen) T.H.: 1S80021N, G.N.: 12C5923N, MatheMorphosis: G0D3420N, and FWO-INSITE: G085018N; by the European Union’s Horizon 2020 Framework Program (H2020/2014-2021) ERC INSITE (772418) and SC1-DTH ISW (101016503); by the European Union’s Horizon Europe Framework Program (HEU/2022-2027) ERC INSTant CARMA (101088919), as well as by the special research fund of the KU Leuven (C24/17/077). The funders had no role in study design, data collection and analysis, the decision to publish, or the preparation of the manuscript. This work is part of Prometheus, the KU Leuven R&D division for skeletal tissue engineering (http://www.kuleuven.be/prometheus).

