## Supplementary figures and images for "An integrated single-cell atlas of the limb skeleton from development through adulthood"

### Supplementary Figure 1

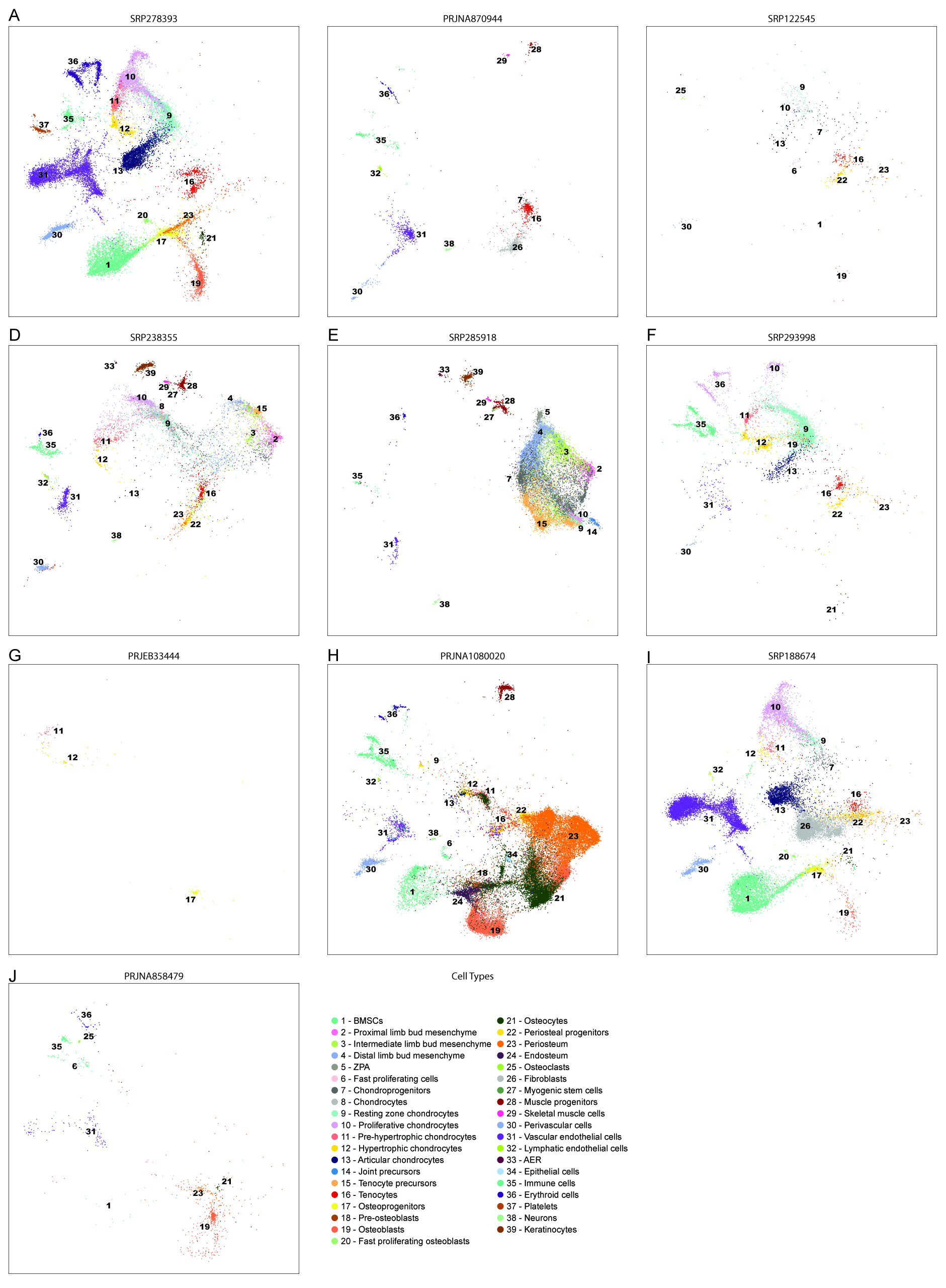

### Supplementary Figure 1

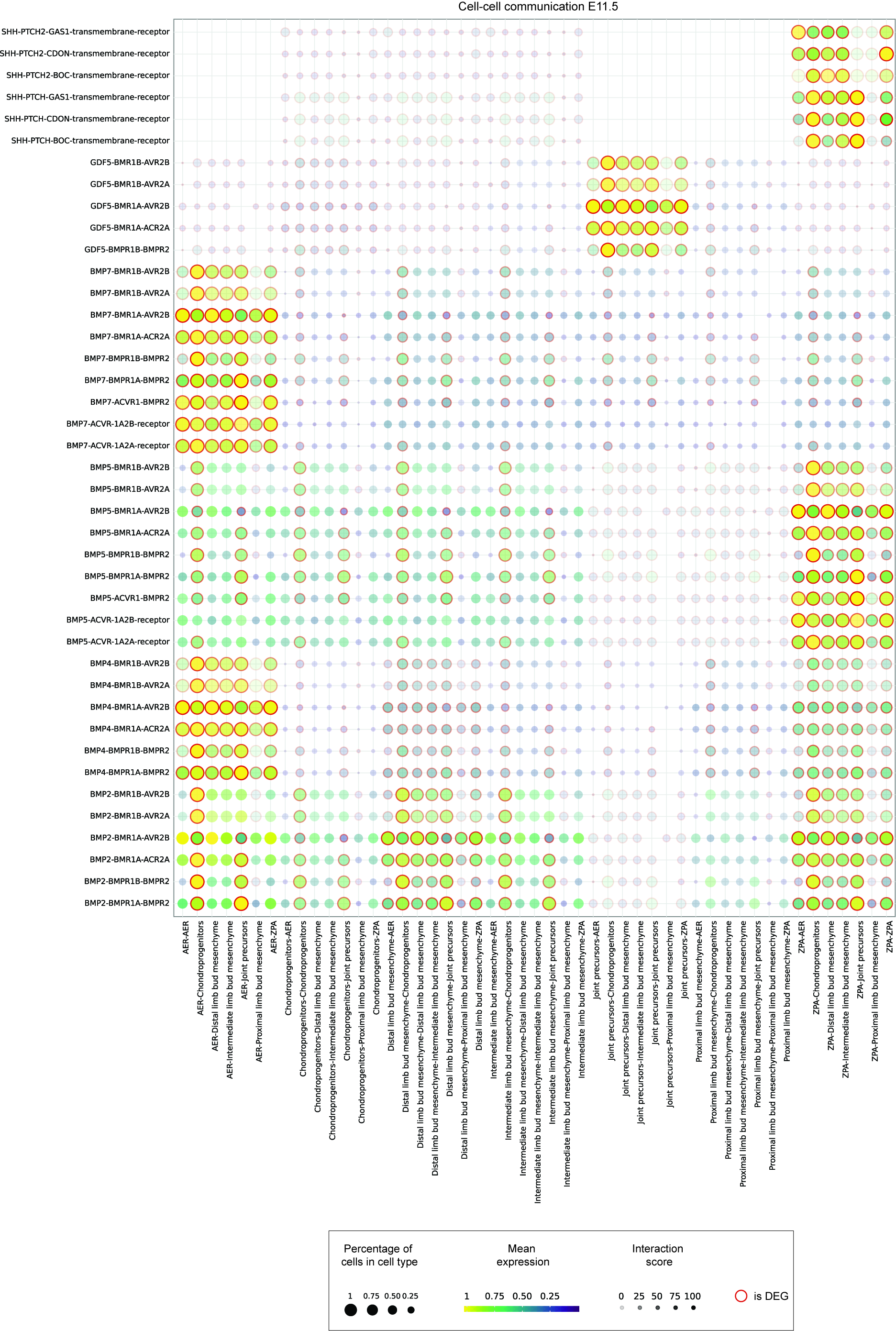

### Supplementary Figure 2

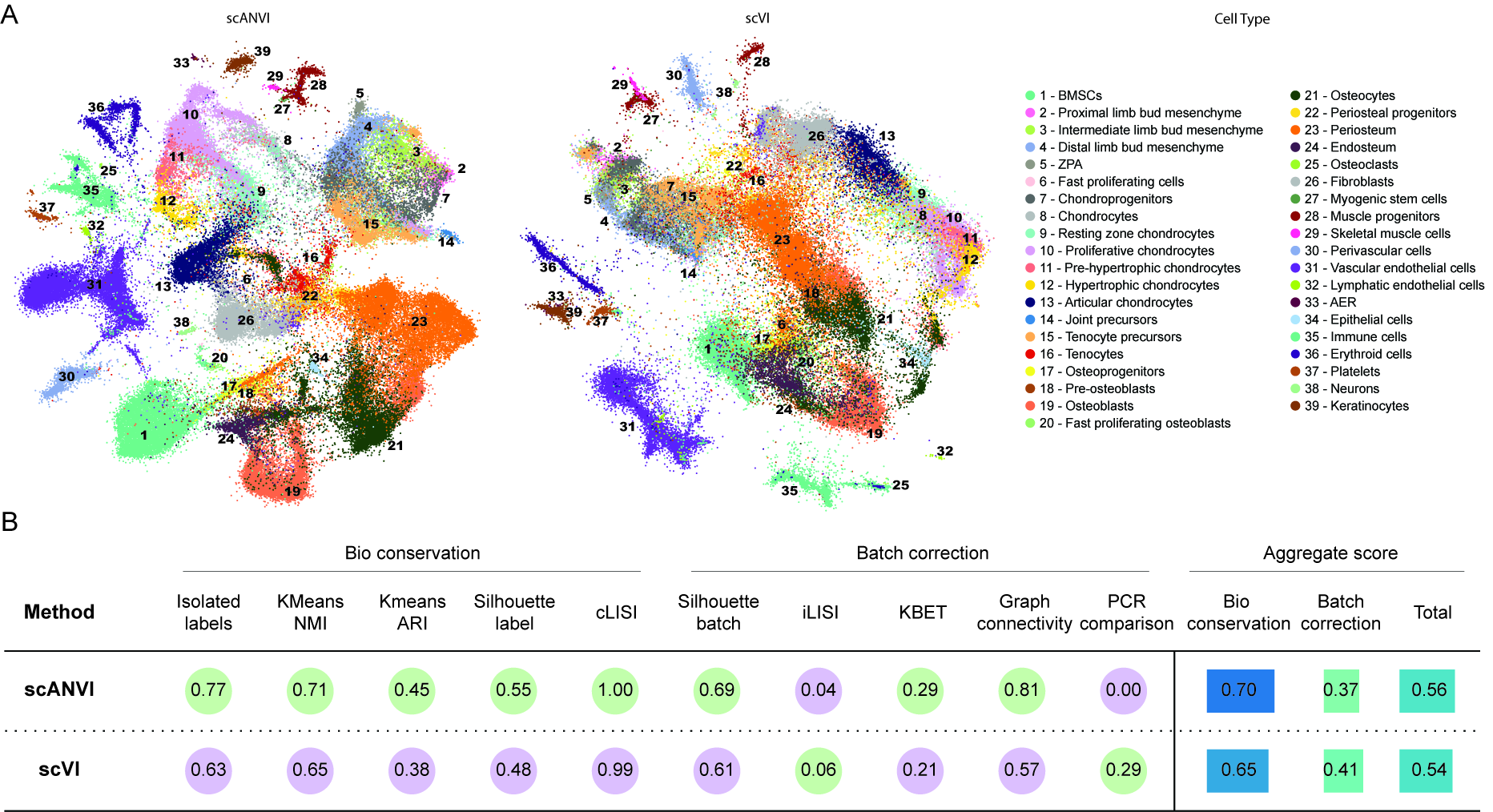

### Supplementary Figure 3

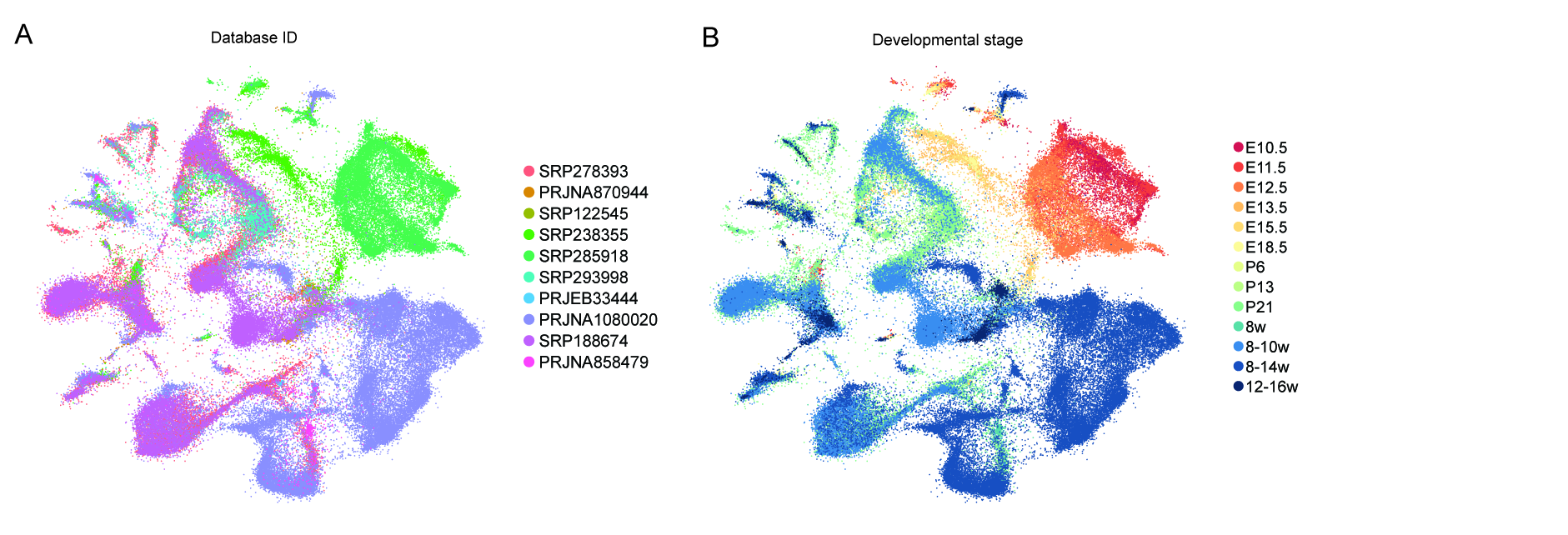

### Supplementary Figure 4

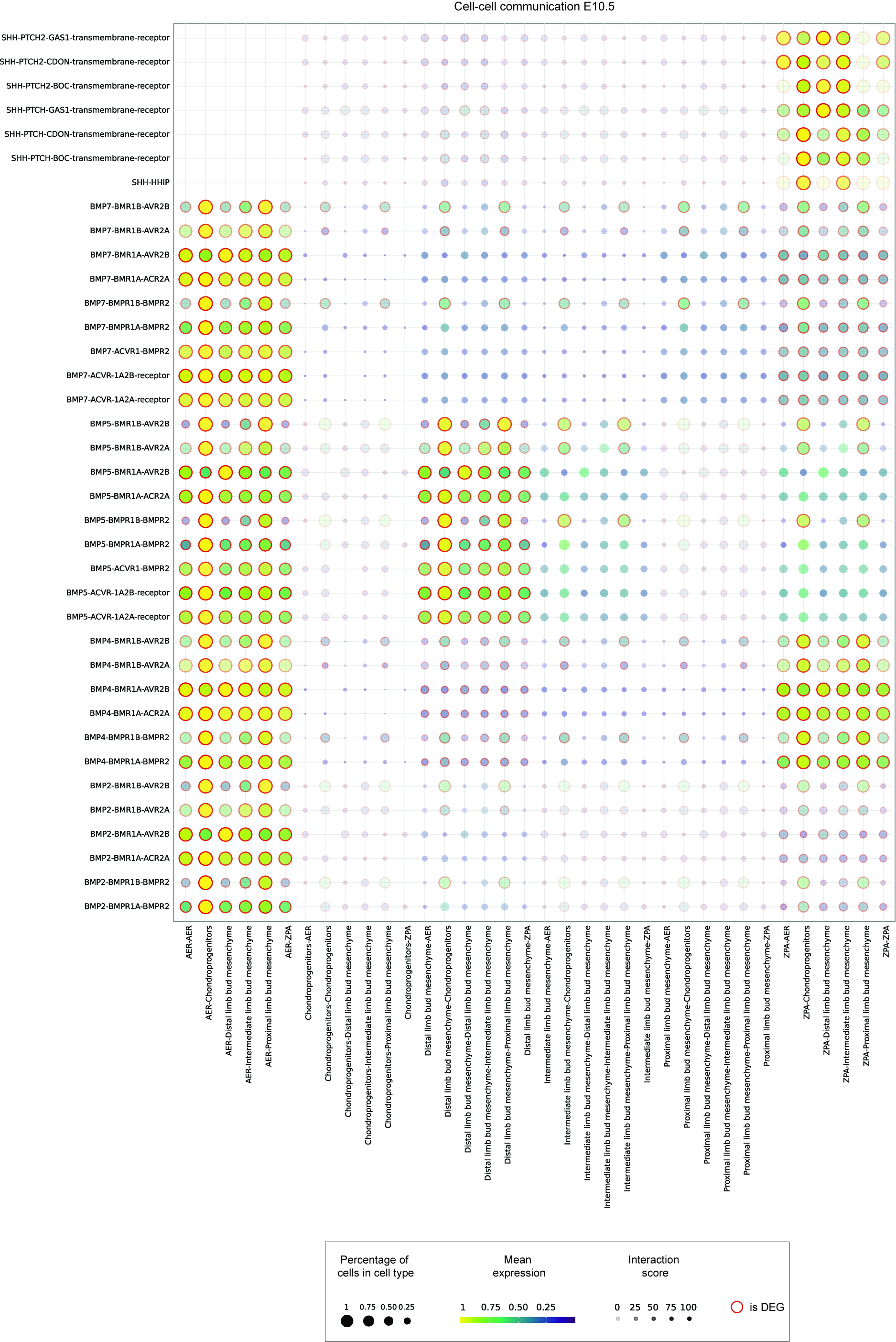

### Supplementary Figure 6

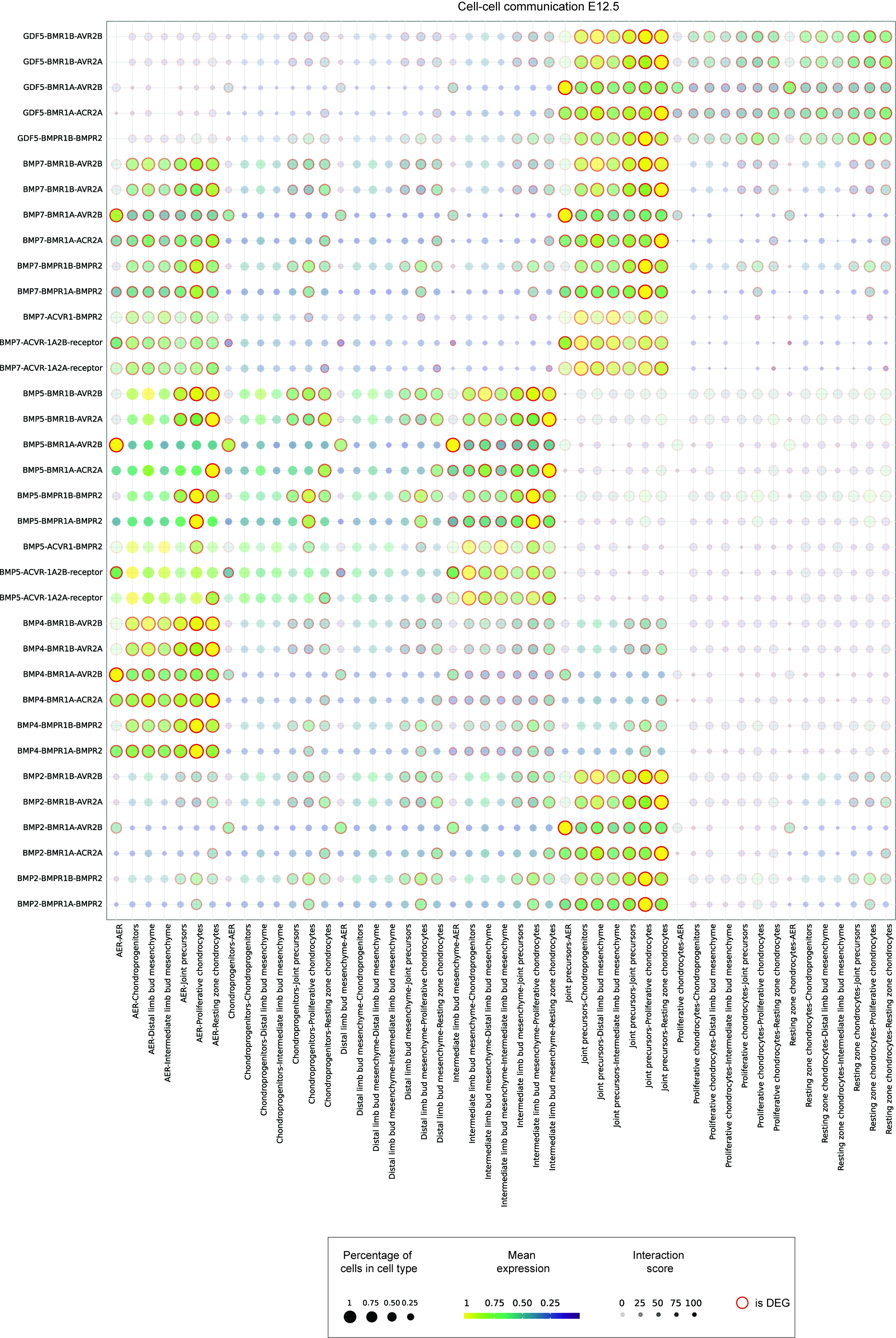

### Supplementary Figure 7

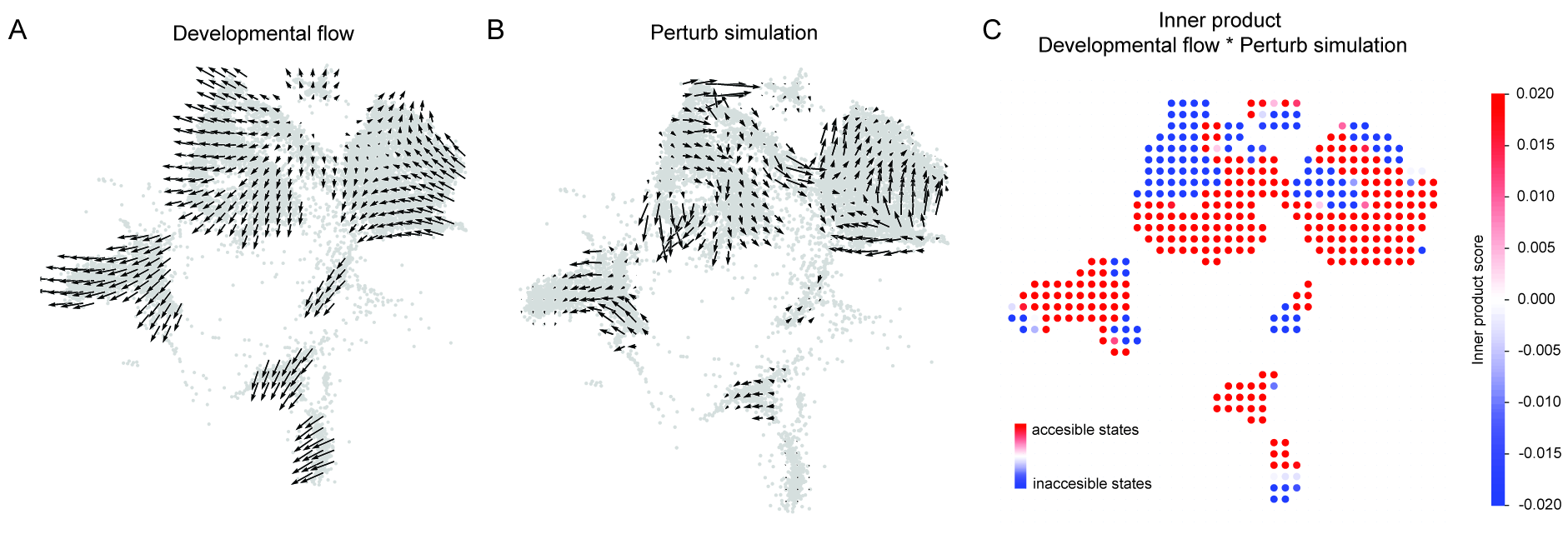
